# Manganese oxide biomineralization is a social trait protecting against nitrite toxicity

**DOI:** 10.1101/294975

**Authors:** Christian Zerfaß, Joseph A. Christie-Oleza, Orkun S. Soyer

**Affiliations:** School of Life Sciences, BBSRC/EPSRC Warwick Integrative Synthetic Biology Centre (WISB), University of Warwick, Coventry, CV4 7AL, UK.; Warwick Integrative Synthetic Biology Centre (WISB), University of Warwick, Coventry, CV4 7AL, UK.

**Author notes:** Corresponding Author (O.S.S.).

**Keywords:** microbial ecology, biomineralization, metal recovery, biotechnology, community function, social behavior, exoenzyme, reactive oxygen species, *Roseoabacter sp.* AzwK-3b

## Abstract

Manganese bio-mineralization by oxidation is a costly but, still, widespread process among bacteria and fungi. While certain potential advantages of manganese oxidation have been suggested, to date there is no conclusive experimental evidence for, how and if this process impacts microbial fitness in the environment. Here we show how a model organism for manganese oxidation, *Roseobacter sp.* AzwK-3b, is growth-inhibited by nitrite, and that this inhibition is mitigated when manganese is added to the culture medium. We show that manganese-mediated mitigation of nitrite-inhibition is dependent on the culture inoculum size, with larger inocula being able to withstand higher concentrations of nitrite stress. Furthermore, the bio-mineralized manganese oxide (MnO_X_) forms granular precipitates in the culture, rather than sheaths around individual cells. These findings support the notion that MnO_X_ is a shared community product that improves the cultures’ survival against nitrite-stress. We show that the mechanistic basis of the MnO_X_ effect involves both its ability to catalyze nitrite oxidation into (non-toxic) nitrate under physiological conditions, and its potential role in influencing redox chemistry around reactive oxygen species (ROS). Taken together, these results provide for the first direct evidence of improved microbial fitness by ^MnO_X_ deposition in an ecological setting, i.e. mitigation of nitrite toxicity, and point to a key role of MnO_X_ in handling stresses arising from ROS. These findings could be of general^ relevance for all organisms oxidizing manganese, allowing them to offset costs associated with extracellular bio-mineralization.

## Introduction

A large variety of biominerals based on different cations (e.g. iron, manganese, calcium) and anions (e.g. carbonates, oxides, phosphates) are deposited by different microorganisms (1). One of these is manganese oxide (2–5), which is deposited by the oxidation of soluble Mn^II^. Microbial Mn^II^ oxidation received attention with the discovery of polymetallic, manganese-rich biogenic deep sea nodules, which have been shown to harbor both manganese-oxidizing, and manganese-reducing organisms (6). While it is suggested that such nodules could potentially be mined for rare earth elements, and that the associated metal-active organisms utilized in biotechnology of metal recovery (2, 3, 5–8), it remains unclear in many cases why organisms show such metal-oxidizing and -reducing activities. In the case of metal-reducing organisms, it has been shown that metabolic energy can be gained under anaerobic conditions from using metal oxides (i.e. manganese, iron, or others) as an alternative terminal electron acceptor (9–11). The potential evolutionary advantages of metal-oxidation, and in particular manganese oxidation meanwhile is not well understood (2, 7, 8).

Some metals can be oxidized by microbes and act as an inorganic energy source for so-called chemolithotrophic growth, as in the case of iron lithotrophy (12). Chemolithotrophy on manganese has been suggested but little experimental evidence has been found so far (2). Two other common hypotheses for manganese oxidation are that the resulting manganese oxides (MnO_X_) can icrease accessibility of organic nutrients or protect microbes from potentially toxic compounds (13). The validity of the latter hypothesis remains to be tested conclusively. MnO_X_ has been shown to react with complex organic (i.e. humic) substances (14), but it is not clear if the resulting organic products form such reactions are utilized by microbes. It is suggested that certain fungi employ ligand-stabilized Mn^III^ to oxidize recalcitrant litter (15), but these studies were not performed with single (defined) carbon/energy sources. Similarly, the former hypothesis regarding the protective potential of MnO_X_ remains unproven to date (2, 7). It has been suggested that MnO_X_ precipitates can act as strong sorbents of heavy metals, hence mitigating the toxic effects of such metals on microorganisms, but this has yet to be tested in a biological context (2). Taken together, the biological significance of microbial manganese oxidation remains a paradox, as no benefits have been demonstrated for this costly metabolic process

In recent years, *Roseobacter sp.* AzwK-3b emerged as a model organism to study the generation of MnO_X_ (16). AzwK-3b is a bacterium that shows significant manganese oxidizing activity *in vitro* when grown in a complex (rich) K-Medium (16). This activity was shown to be mediated by a secreted exoenzyme - a haem type oxidase - that can catalyze the generation of superoxides from NADH and oxygen. The resulting superoxide can in turn facilitate the Mn^II^ oxidation into Mn^III^, which undergoes further disproportionation to result in MnO_2_ (17–21) – or more specifically mixed valence state MnO_X_. The required NADH for this exoenzyme-mediated reaction is presumably secreted also by AzwK-3b (17). Thus, these mechanistic findings strongly suggest that AzwK-3b is making a significant metabolic investment into production of MnO_X_. It is currently not clear how such a costly strategy benefits individual cells and how it could have been maintained over evolutionary timescales.

In an attempt to better understand the ‘fitness’ impact of manganese oxidation, we have studied the physiology of *Roseobacter sp.* AzwK-3b in more detail. We identified a defined medium composition that allowed growth of this bacterium both with and without manganese. While we found no significant differences in growth rate under these two conditions, we found that the manganese oxidizing activity of *Roseobacter sp.* AzwK-3b supports growth of the bacterium at nitrite concentrations that fully prevent growth in a manganese-free culture. We found that MnO_X_ forms as granules dispersed among cells, and its nitrite-inhibition mitigation effects show a significant population size effect, conforming to a ‘community commodity’ nature of this compound. Mechanistically, we show that biogenic MnO_X_ was able to catalyze nitrite oxidation into nitrate under physiological conditions, and that the mitigation of nitrite-inhibition was also affected by NADH. These results suggest that the ability of MnOx to alleviate nitrite toxicity relates to providing catalytic scavenging of reactive oxygen species (ROS) within the environment.

## Results

To study the role of manganese oxidation on microbial fitness we have focused here on *Roseobacter sp.* AzwK-3b, which has recently emerged as a model organism for this process (2, 8). We refer to the oxidation product as MnO_X_, since biogenic manganese oxides are usually precipitates with mixed manganese oxidation states, particularly Mn^II^, Mn^III^ and Mn^IV^ (2, 26). AzwK-3b has been shown to oxidize manganese to MnO_X_ by means of an excreted exoenzyme and NADH, and potentially involving an elaborate redox reaction path (17–21). We have first attempted to identify fully-defined growth conditions for this bacterium, which has been to date studied in complex K-medium (16), an artificial seawater derived, peptone/yeast extract containing medium (16, 27). Through systematic analysis of media composition, we have created a fully defined medium that supports AzwK-3b growth (from now on referred to as modified artificial seawater medium, ASW_m_) (Table 1). This exercise revealed also the requirement for five vitamin supplements for growth (Figure S1). Given this defined culture medium, we were then able to interrogate the impact of manganese on the growth of AzwK-3b.

**Table 1.** Detailed composition of the defined AzwK-3b growth medium, ASW_m_. The medium was developed starting out from artificial seawater (ASW) (22) with extra trace metals taken from (9, 81) and a 5-vitamin solution identified starting out from Wolfe’s vitamin mixture (82).

|  |  |
| --- | --- |
| **<u>Base salts (1 x AzwK-3b medium)</u>** |  |
| Sodium chloride (NaCl) | 200 mM |
| Ammonium chloride (NH<sub>4</sub>Cl) | 8.82 mM |
| Potassium chloride (KCl) | 6.71 mM |
| di-potassium hydrogenphosphate (KH<sub>2</sub>PO<sub>4</sub>) | 131 µM |
| Magnesium sulphate (MgSO<sub>4</sub>) | 14.2 mM |
| Magnesium chloride (MgCl<sub>2</sub>) | 9.84 mM |
| Calcium chloride (CaCl<sub>2</sub>) | 3 mM |
| Tris(hydroxymethyl)aminomethane (TRIS) | 1.1 mM |
| **pH of the medium** | **8.0** |
| **<u>Trace metal solutio (1,000 x)</u>** |  |
| Copper chloride (CuCl<sub>2</sub>) | 32 µM |
| Zink sulphate (ZnSO<sub>4</sub>) | 765 µM |
| Cobalt chloride (CoCl<sub>2</sub>) | 169 µM |
| Sodium molybdate (Na<sub>2</sub>MoO<sub>4</sub>) | 1.65 mM |
| Boric acid (H<sub>3</sub>BO<sub>3</sub>) | 46.3 mM |
| Nickel chloride (NiCl<sub>2</sub>) | 4.2 mM |
| Sodium tungstate (Na<sub>2</sub>WO<sub>4</sub>) | 243 µM |
| Sodium selenite (Na<sub>2</sub>SeO<sub>3</sub>) | 228 µM |
| **<u>Additional (1,000 x) supplement solutions</u>** |  |
| Iron chloride (FeCl<sub>3</sub>; prepared in 10 mM HCl, balanced with extra 10 mM NaOH solution) | 10.4 mM |
| Ethylenediaminetetraacetate (EDTA, pH 8.0; sodium salt) | 1.34 mM |
| Manganese chloride (MnCl<sub>2</sub>, only added where desired) | 200 mM |
| **<u>Vitamin supplement (1,000 x)</u>** |  |
| Biotin | 82 µM |
| Pyridoxine hydrochloride | 484 µM |
| Thiamine hydrochloride | 148 µM |
| Riboflavin | 133 µM |
| Nicotinic acid | 406 µM |

### Manganese oxidation does not impact growth rate

Despite potentially significant costs associated with exoenzyme and NADH investment, we did not find any substantial difference in growth rates and steady state population sizes with increasing Mn^II^ concentration and with 25 mM acetate (Figure 1). A slightly slower growth at the highest manganese concentration (500 μM) was observed, but it was difficult to ascertain this effect, as both MnO_X_ particles and cells co-aggregating with those particles could have interfered with the absorbance measurements. The slightly reduced growth rate at 200 μM Mn^II^Cl_2_ is in line with an earlier report on AzwK-3b, where 100 μM Mn^II^ was found to decrease the growth rate in (complex) K-medium (16). Other manganese-oxidizing bacteria, such as *Erythrobacter sp.* SD-21 (28, 29) and a marine *Bacillus* strain (30), were reported to grow better when cultured with Mn^II^-supplement. In light of these different findings and possible difficulties with growth rate measurements in the presence of manganese precipitation, we cannot be fully conclusive about the growth effects associated with manganese oxidation based on the presented results, however, they are suggestive of a low or no-impact on growth rate.

**Figure 1.**
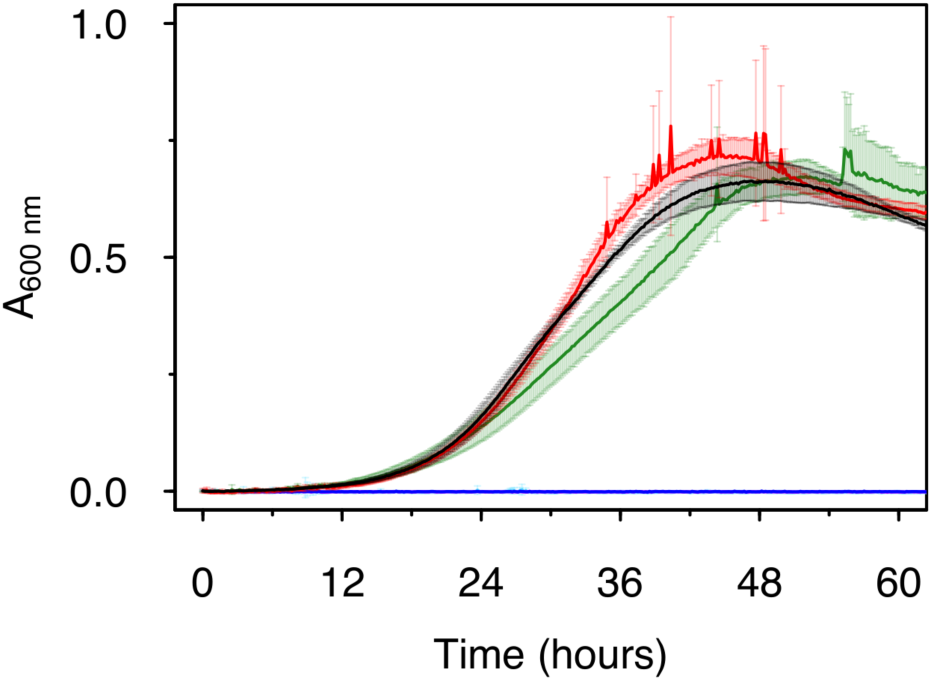
Effect of Mn^II^ on the growth of *Roseobacter sp.* AzwK-3b in the defined growth medium (see Table 1). The concentrations of manganese were 0 μM (black), 200 μM (red) and 500 μM (dark green), with no growth (zero line) in the respective non-inoculated controls (blue, magenta, light blue). Cultures were grown in a 96 well plate (200 μl culture) with shaking and absorbance measurement every 10 minutes (see *Methods*).

### Manganese oxidation mitigates nitrite growth inhibition

With growth effects being limited, a possible alternative explanation for a positive role of manganese oxidation is a protective effect against inhibitors or stresses (2, 13). Here, we evaluated this hypothesis for nitrite. Nitrite is commonly found in the environment, where it results from the reduction of nitrate, a key terminal electron acceptor utilized by many microbes (31). We found nitrite inhibited the growth of AzwK-3b in manganese-free cultures, where already as little as 0.25 mM nitrite prevented growth of AzwK-3b (Figure 2A). No growth was detected at and above 0.5 mM nitrite. Note that a salinity effect at such low concentrations of nitrite (which was added as sodium nitrite) is highly unlikely. To further rule out this possibility, we additionally analyzed the growth of AzwK-3b at different salinity levels using concentrations of sodium chloride from 200 mM (default in the defined ASW_m_ medium employed here) up to 428 mM (default in the ASW medium (22)). This confirmed that salinity effects on growth in this range are minimal (Figure S2), and higher salinity is rather favorable for AzwK-3b growth. Thus, the effects of nitrite are due to toxicity rather than salinity.

**Figure 2.**
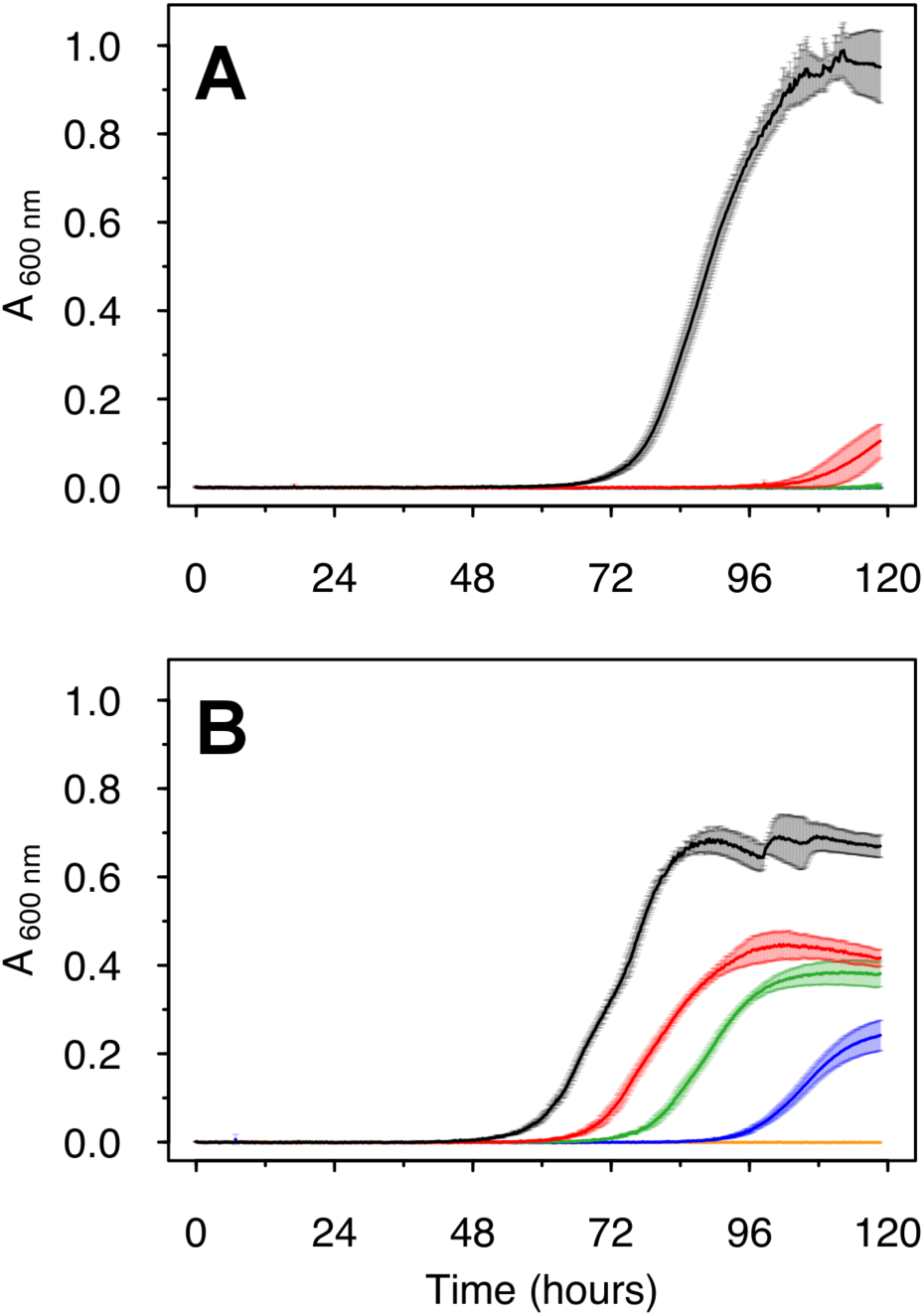
Growth of *Roseobacter sp.* AzwK-3b in the defined growth medium supplemented with sodium nitrite. Media were prepared without (Figure A) or with (Figure B) 200 μM manganese chloride, Mn^II^Cl_2_. Nitrite-concentrations were 0 mM (black), 0.25 mM (red), 0.5 mM (green), 1 mM (dark blue) and 2.5 mM (light blue). All conditions were tested in triplicates, and the growth curves represent averages and their standard deviations (see *Methods*).

With the addition of 200 μM Mn^II^, we found that AzwK-3b is able to grow in the presence of up to 1 mM nitrite (Figure 2B). Increasing the nitrite concentration still affected both the growth rate and maximal culture density (based on A_600_), but this effect was much lower compared to the manganese-free cultures (Figure 2). To overcome any potential confounding effects of MnO_X_ precipitation on spectroscopic culture density measurements, we additionally quantified acetate consumption by ion chromatography as a proxy for growth. As expected, manganese-free cultures with 0.25 mM (or higher) nitrite showed only insignificant decrease in acetate, while the Mn^II^ supplemented cultures showed acetate consumption in accordance with the A_600_ measurements (see Figure S3). These findings confirm that Mn^II^ supplementation allows AzwK-3b to withstand nitrite inhibition.

### Nitrite-inhibition relief is a community function that depends on culture size and that is mediated by dispersed, granular MnO_X_ precipitates

It has been shown that MnO_X_ precipitation by AzwK-3b is mediated by secreted exoenzymes (17). It is not known, however, whether the process of MnO_X_ precipitation occurs primarily on individual cell surfaces, or whether it is a population level process with the secreted enzymes conferring to the notion of a “community commodity” (32–35). We hypothesized that these two different scenarios could be distinguished by analyzing population size effects on MnO_X_ mediated mitigation of nitrite-inhibition. In particular, we designed an experiment in which cultures pre-grown without Mn^II^ are subsequently sub-cultured into media with Mn^II^ and nitrite, using different inoculum size (Figure S4). We argue that in the case of MnO_X_ precipitation being a process confined to individual cells, there should be no effect of inoculation size.

We found that manganese mediated mitigation of nitrite inhibition was dependent on inoculum size (Figure 3). A pre-culture was grown without nitrite and manganese, and from this, inocula were generated at two different time points within the first third of the exponential phase (labelled IT1 and IT2 in Figure S4). When these inocula were subjected to nitrite in the main-culture, the earlier, low-density inoculum IT1 was inhibited by nitrite regardless of the presence or absence of Mn^II^ (Figure 3 A,B), while manganese-mediated mitigation of nitrite inhibition was clearly evident for the larger, high-density inoculum IT2 (Figure 3 C,D). In the IT1 cultures half of the acetate was unused at 0.25 mM nitrite, and gradually more acetate resided with increasing nitrite concentration (Figure S5). In the IT2 cultures with Mn^II^ supplementation, however, acetate was completely removed at all nitrite levels below 2.5 mM and only 25 – 50 % of acetate remained at 5 – 10 mM nitrite. In the control samples (no inoculation) there was no change in acetate concentration ruling out any cross-activity with manganese.

**Figure 3.**
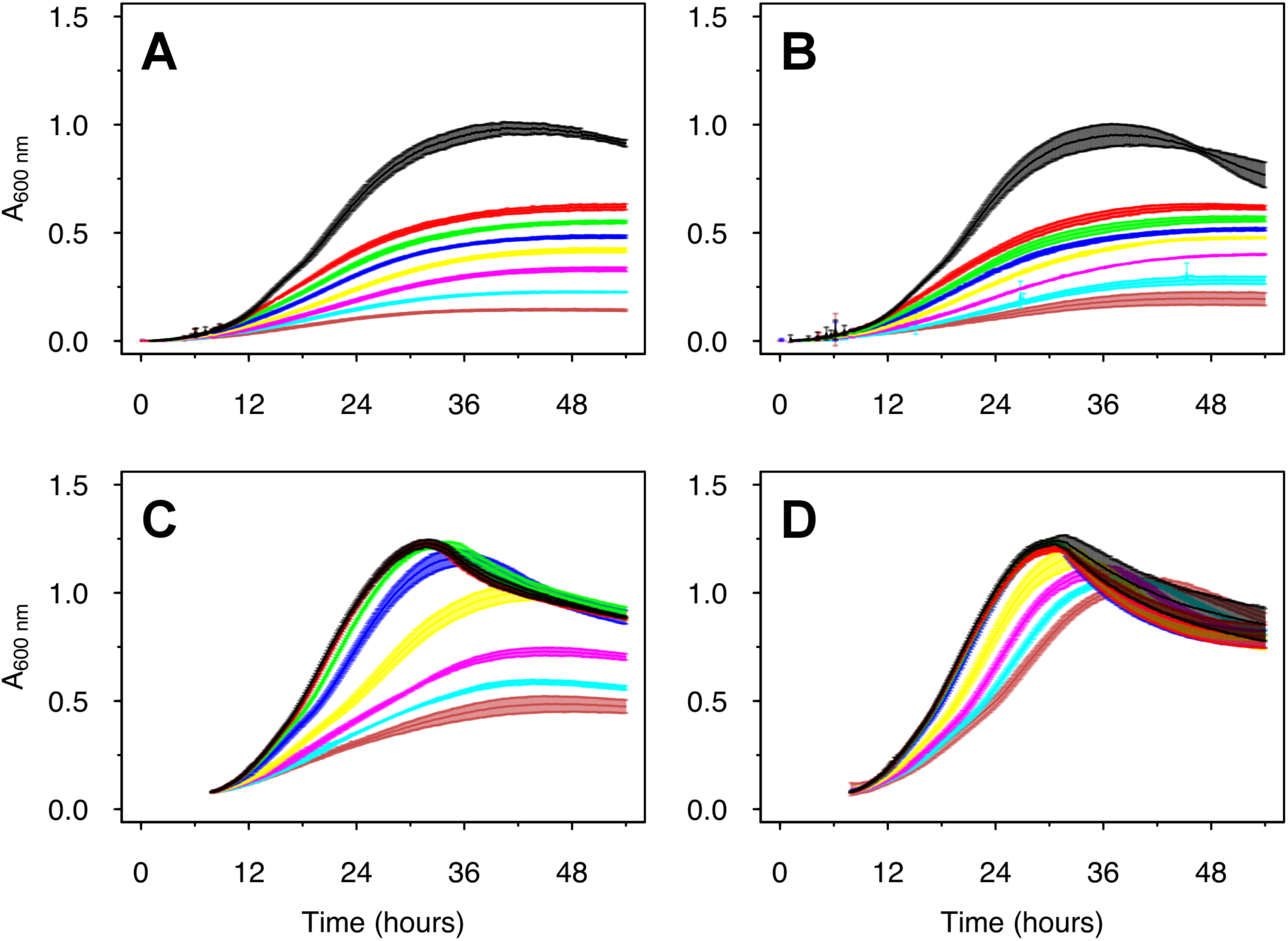
Larger AzwK-3b inocula are less inhibited by nitrite. A pre-culture without manganese or nitrite was grown and sampled in the exponential growth phase (Figure S4) to prepare inocula from a very early time point in the exponential phase (IT 1, Figures A and B), and from a later time point (IT 2, Figures C and D; both sampled in first third of exponential phase). These inocula were 1:1 diluted with fresh medium, and tested for growth at different nitrite concentrations (see below for colour code) without (A, C) or with (B, D) 200 μM MnIICl2 supplement. The nitrite concentrations were: Black – control no nitrite. Red – 0.25 mM nitrite. Green – 0.5 mM nitrite. Blue – 1 mM nitrite. Yellow – 2 mM nitrite. Magenta – 5 mM nitrite. Light blue – 7.5 mM nitrite. Dark red – 10 mM nitrite. Growth curves show the averages and standard deviations over a triplicate analysis (see *Methods*).

Rather than a true population size effect, these observed inocula effects could be due to cells from the Mn-free, early-phase pre-cultures not having ‘turned on’ expression of exoenzymes required for MnO_X_ precipitation. To rule out this possibility, we performed an additional experiment, where the pre-cultures were already grown with 200 μM Mn^II^. Using this pre-adapted culture, inocula were again prepared by sampling at different growth time points (IT 1 – 4 in Figure S6, A). Cultures grown from these different inocula displayed much weaker inhibition by increasing nitrite concentrations up to 10 mM (Figure S6, B) and were able to consume acetate (Figure S6, C), yet there were still inoculum size effects on overcoming nitrite inhibition (Figure 4, green). Interestingly, the extent of this effect seems similar to that observed with inocula originating from pre-cultures grown without Mn^II^ but supplied with Mn^II^ after subculturing into nitrite containing media (Figure 4, blue). In particular, at 5 and 10 mM nitrite, maximum growth rate (and final density) data from all these cultures showed a strong nonlinear correlation to initial inocula density that can be fitted to a sigmoidal curve. (Figure 4, black line). The infliction point of this curve happened at a lower inoculum size for those cultures that were not supplied with Mn^II^ at any stage (Figure 4, red). Thus, we conclude that there is an inoculum density effect on the ability of Mn^II^ supplemented cultures to tolerate nitrite irrespective of their culturing history, but that this effect is stronger for cultures not pre-grown with Mn^II^. There were no such effects without nitrite or without Mn^II^ (Figure 4).

**Figure 4.**
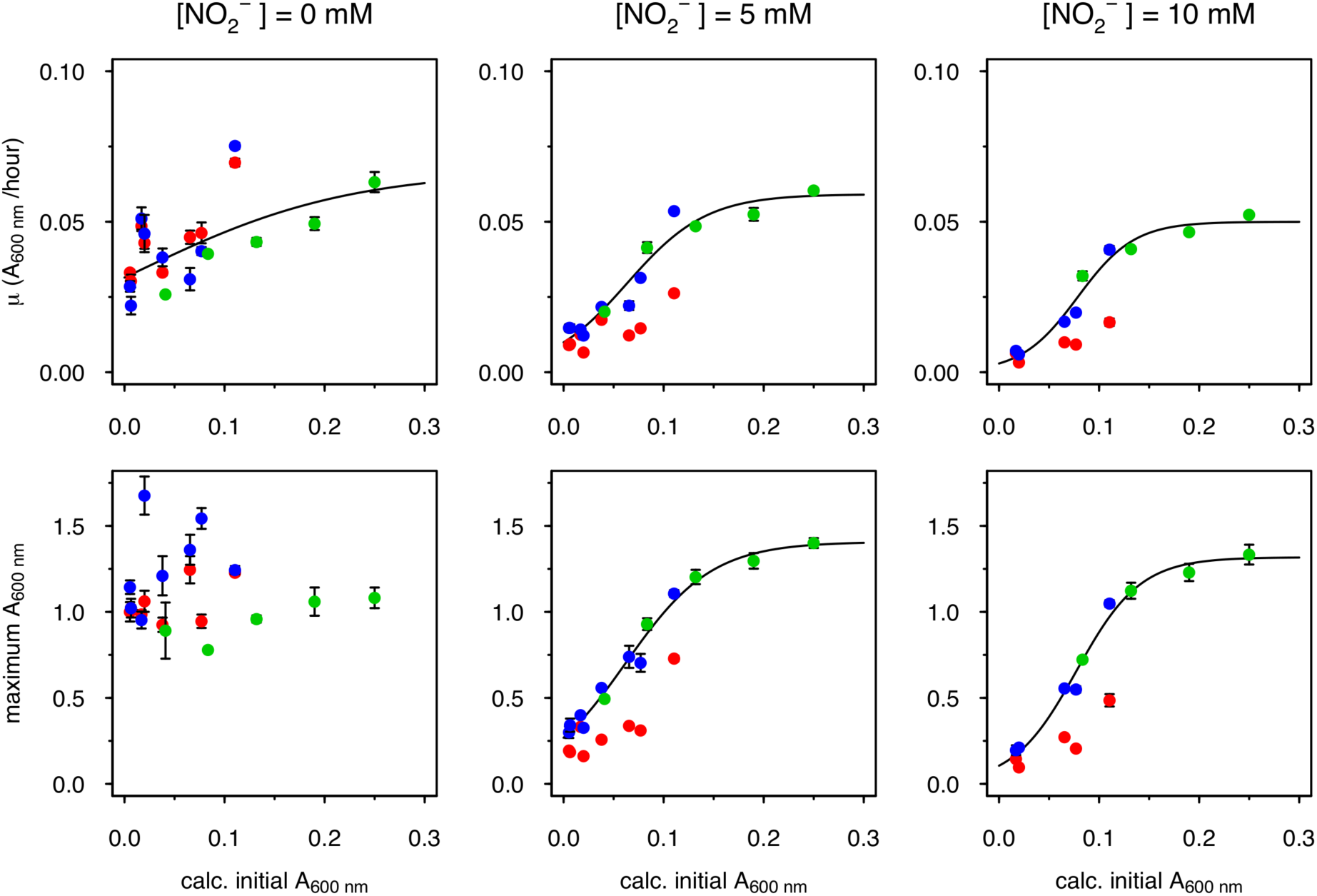
Inoculum-size effect on MnO_X_ mediated mitigation of nitrite-inhibition. Data from different AzwK-3b growth experiments of similar type (“Large inocula”, see *Methods*) were analyzed for the maximum A_600_ and growth rate by fitting the growth curves. Nitrite-concentrations of the main-cultures are indicated as headings of the figure-rows. The x-axes show the calculated A_600_ of the cultures after diluting them 1:1 for the main-culture, while the y-axes show the maximum A_600_ and maximum growth rate as calculated with the Gompertz model (23, 24)) (see *Methods*). The colours represent different conditions: **Red:** Neither pre-, nor main-culture contained manganese; **Blue:** Pre-culture without, main-culture with manganese; **Green:** both pre- and main-culture with manganese. The black curve is a sigmoidal fit (logistic model) from the Grofit R-package (23), for the results of the combined blue and green dataset where the nitrite-exposed main-cultures all contained manganese.

These results strongly suggest that MnO_X_ precipitation is a community level function. To further collaborate on this result, we explored the micro-structure of the AzwK-3b cultures in the presence of Mn^II^. Analysis of cultures using electron microscopy revealed that MnO_X_ precipitates as granules dispersed within the culture, and attaching to clusters of cells, rather than forming sheaths around individual cells (as seen in some other cases of metal oxide precipitations (36)) (Figure 5, left). Employing electron dispersive X-ray spectroscopy, we confirmed that these granular structures contained manganese, while no manganese was detected in locations with cells only (i.e. without granular structures, see Figure 5, right).

**Figure 5.**
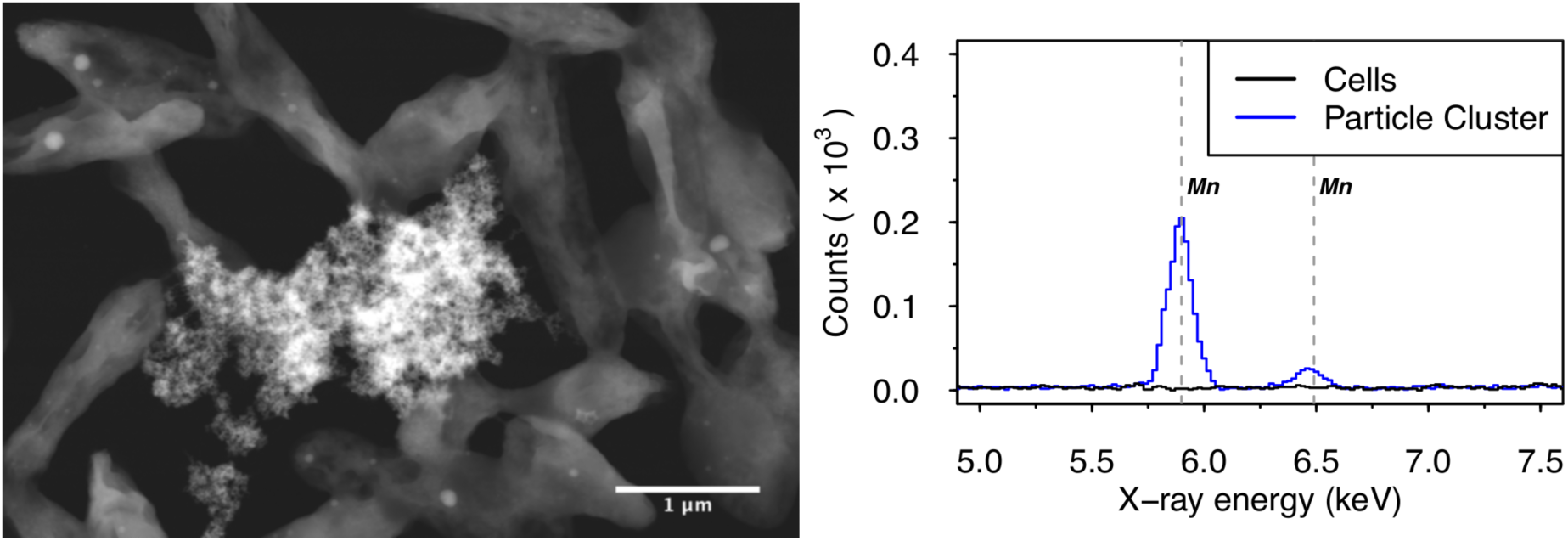
Scanning transmission electron micrograph (left figure, high angle annular dark field) of (granular) manganese-containing precipitate (center) surrounded by AzwK-3b cells, and associated energy dispersive X-ray spectroscopic analysis (right figure) in this location. Only the energy range containing the manganese-specific X-ray energies at 5.90 keV (K_α_^I^) and 6.49 keV (K_β_^I^) is shown, and the manganese transitions are indicated by vertical gray dashed lines.

### MnO_X_ mediated nitrite protection involves redox reactions and oxygen radicals

After establishing the community level functionality of biogenic MnO_X_ as a protective agent against nitrite, we next wanted to evaluate the mechanistic basis of this function in the context of nitrite toxicity. While multiple mechanisms of nitrite-toxicity are reported (37, 38), two key reactive species are usually implicated, i.e. free nitrous acid (39) and peroxynitrite. The former forms through protanation of nitrite, while the latter forms from the reaction of nitrite with hydrogen peroxide (40–42). Thus, two non-exclusive, possible mechanisms of MnO_X_ relief on nitrite toxicity are: (i) MnO_X_ catalyzed oxidation of nitrite to nitrate (a reaction that has been shown to be feasible chemically under low pH (43)) and thereby avoiding formation of either free nitrous acid or peroxynitrite; or (ii) MnO_X_ catalyzed degradation of hydrogen peroxide and thereby avoiding the reaction of this compound with nitrite to form peroxynitrite.

To see if AzwK-3b generated MnO_X_ can catalyze nitrite oxidation under physiological conditions, we collected it from culture supernatants and evaluated its reactivity with nitrite in our ASW_m_-medium at pH = 8.0. Over 27 days, we found nitrite oxidation by biogenic MnO_X_ in a dose dependent manner, while neither synthetic MnO_2_ powder nor the MnO_X_-free supernatant solution showed any significant nitrite oxidation (Figure 6A). The trend of nitrite oxidation matched with nitrate production (Figure 6B), thus confirming the assumed reaction pathway of nitrite-oxidation into nitrate (43). Taking into account the difficulties of accurately determining the amount of precipitated MnO_X_ that were added into the nitrite assay, we can still estimate that the condition with highest MnO_X_ levels contained at least 1-2 mM (with respect to Mn). This presents a stoichiometric minimum 2-fold excess over nitrite (at 0.5 mM), hence enough for complete nitrite oxidation. The fact that this reaction didn’t proceed further than an oxidation of ~0.18 mM nitrite (i.e. ~35 %) indicates that either the biogenic MnO_X_ was only partially reactive or that its reactivity reduced over time (as known to be the case for synthetic manganese oxides (2, 13)). Sample pH remained relatively stable with the biogenic MnO_X_, while samples without manganese and with synthetic MnO_2_ reached a pH of 6.9 and 6.8, respectively at the end of the experiment (from an initial pH of 8.2 of the medium). This acidification of the control samples might be due to carbon dioxide dissolution, which might have been buffered in the samples with biogenic MnO_X_ due to proton consumption during nitrite oxidation, or due to co-precipitated organic solutes (polymers, proteins) from the cell-free supernatant.

**Figure 6.**
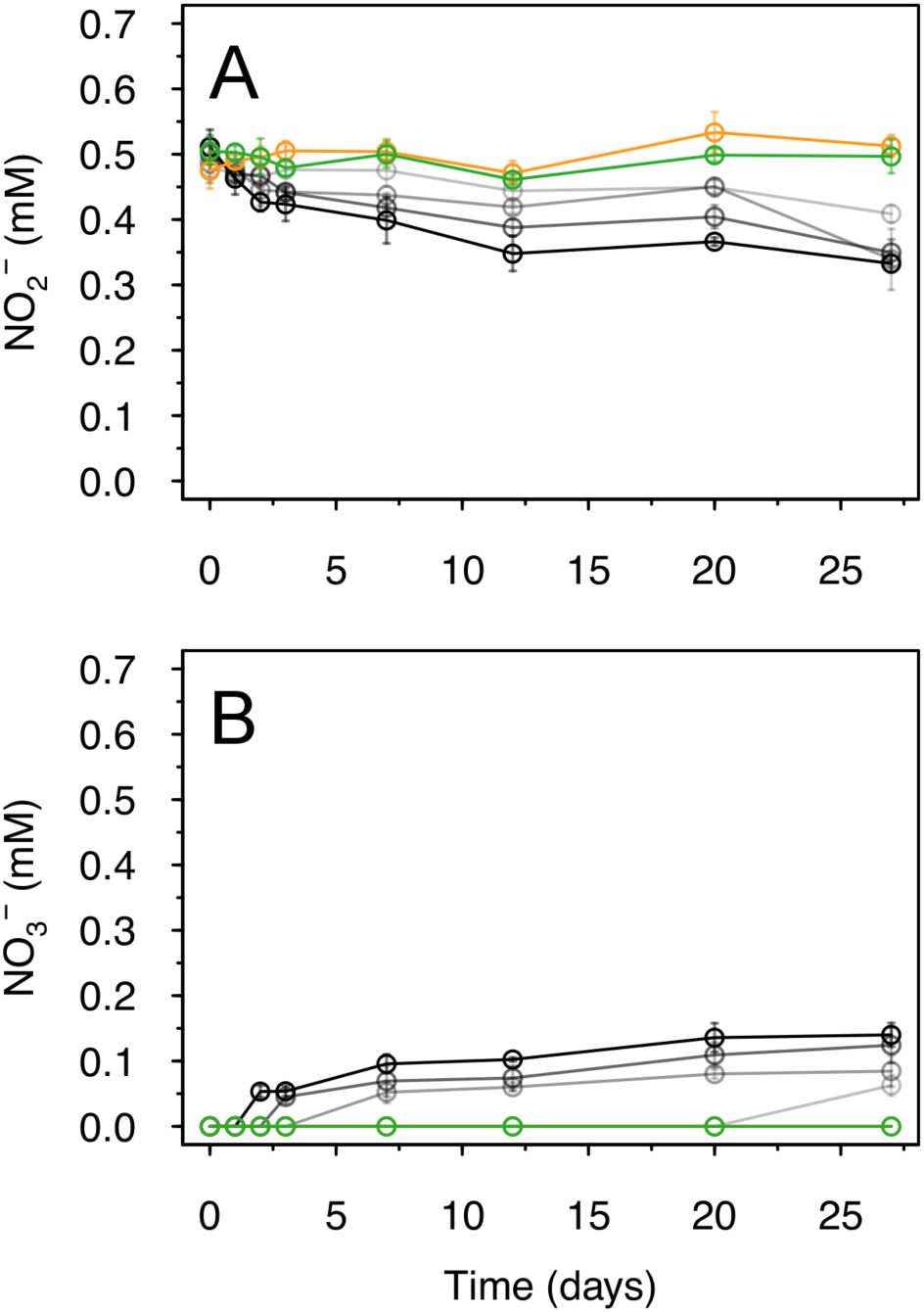
Oxidation of nitrite by biogenic manganese oxide (MnO_X_) produced in cell-free culture supernatant of AzwK-3b. The figures show the concentration of nitrite (A) and nitrate (B), determined by ion chromatography, over time (note that concentrations were corrected for the IC-peak from chloride, to account for evaporation during the experiment). As controls, samples without MnO_X_ (green), or with MnO_2_ powder (orange) were included in the experiment (see *Methods*). The samples with AzwK-3b cell-free manganese oxide contained (from grey to black) 0.2, 0.5, 1 and 2 mM manganese oxide equivalent (see *Methods*).

These findings confirmed that the biogenic MnO_X_ were capable to oxidize nitrite at physiological conditions, and prompted us to test MnO_X_ mediated nitrite oxidation directly in AzwK-3b cultures. We found some evidence for decreasing nitrite concentration in different cultures tested, but this was not significant (Figure S7), and some decrease was also seen in the manganese free cultures (indicating possible measurement effects in the solution). If nitrite oxidation was the main mechanism of MnO_X_ mediated protection *in vivo*, these cultures would have been expected to oxidize most of the nitrite present in the media. Thus, we conclude that under our experiment conditions nitrite-oxidation was only a potential contributing factor.

A plausible alternative mechanism of MnO_x_ mediated nitrite-inhibition relief could be related to formation of reactive peroxynitrite, which is shown to be highly toxic to bacteria (41, 42, 44, 45), and which can form at low pH from the reaction of hydrogen peroxide with nitrite (40). If peroxynitrite is the main species underpinning nitrite toxicity, then, MnO_X_ protection against nitrite could be due to its ability to degrade hydrogen peroxide and thereby reducing the rate of peroxynitrite formation. The reactivity of MnO_X_ towards hydrogen peroxide has been demonstrated chemically (40, 46–53), but never shown or tested in a biological context. Here, we hypothesized that if these types of redox reactions were involved in MnO_X_ mediated mitigation of nitrite-inhibition, the process dynamics can be modulated with the introduction of additional hydrogen peroxide or NADH (which can help increase the rate of MnO_X_ formation (18), but which can also be directly involved in hydrogen peroxide reduction through peroxidase-catalysed reactions (54–58)). To test this hypothesis, we again grew pre-cultures of AzwK-3b without Mn^II^ and sub-cultured these in medium containing Mn^II^ and nitrite, but at the same time also spiking in hydrogen peroxide or NADH. Hydrogen peroxide spiking did not show any effect on nitrite inhibition or its release by Mn^II^ supplementation (Figure S8), possibly due to spiked hydrogen peroxide being cleared primarily through additional peroxidases rather than impacting MnO_X_ mediated process dynamics. In line with this hypothesis, spiking NADH resulted in full mitigation of nitrite inhibitory effect (even without Mn^II^) (Figure 7). This suggests that nitrite toxicity relates to peroxynitrite formation via hydrogen peroxide, which can be decomposed by MnO_X_ (as shown before (40, 46–53)) or NADH-utilizing peroxidases (that are shown to be present in Roseobacter species including AzwK-3b (17, 59) (see also Table S1)).

**Figure 7.**
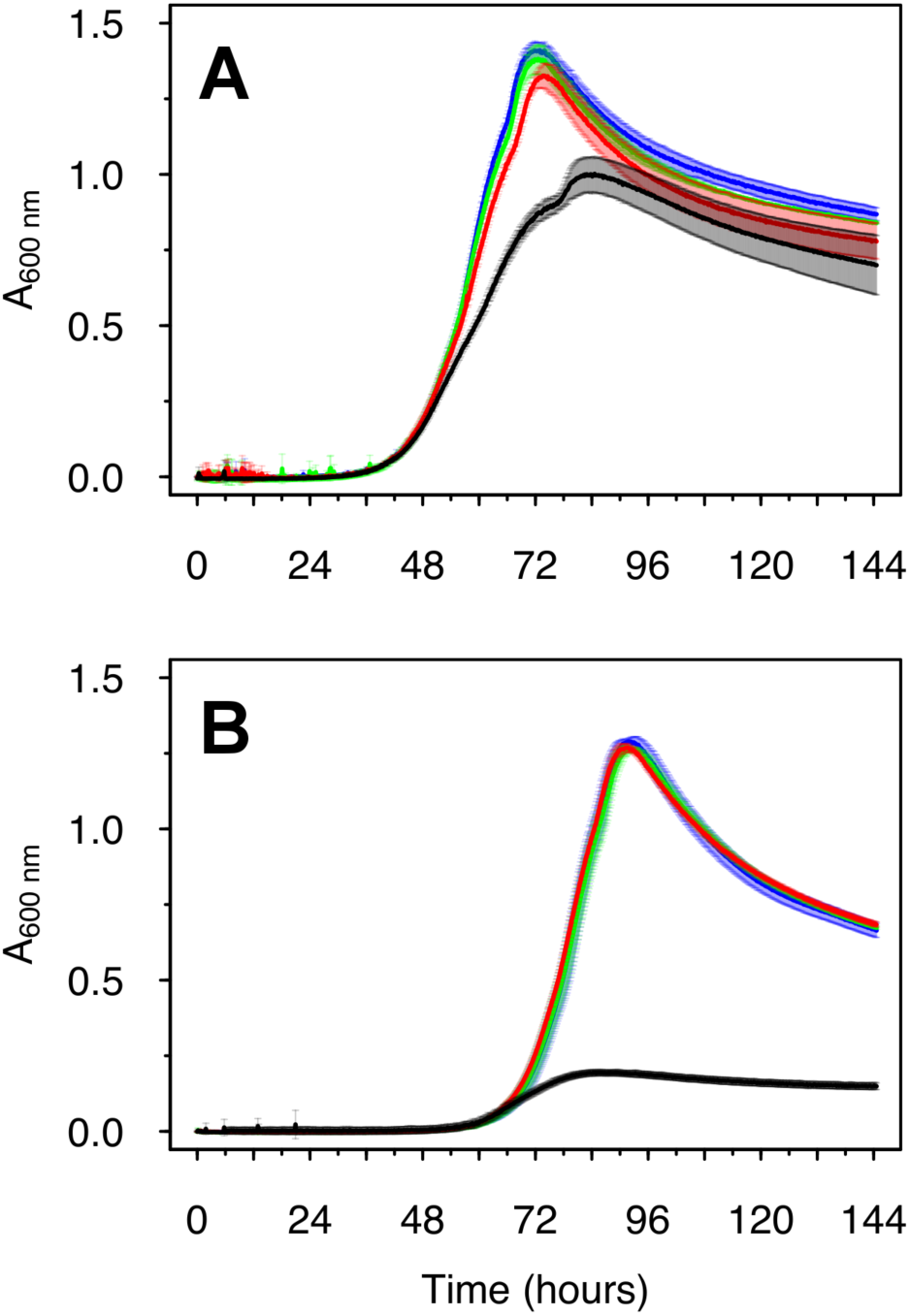
Reductive power (NADH) mitigates the growth inhibitory effects of nitrite in AzwK-3b. Cultures (pre- and main-culture without manganese) were grown in the absence (A) and presence (B) of 5 mM nitrite and supplement of 0, 50, 100 and 200 μM NADH (black, red, green and blue) at the start of the culture.

## Discussion

Manganese bio-mineralization into MnO_X_ is widespread among bacteria, but there is no clarity about its evolutionary advantage. Here, we developed a defined growth media for the manganese oxidizing model organism *Roseobacter sp.* AzwK-3b and demonstrated that this organism’s strong growth-inhibition by nitrite is mitigated through its ability to precipitate biogenic MnO_X_. We found that this MnO_X_-mediated mitigation of nitrite toxicity is dependent on population size, and that MnO_X_ forms dispersed granules that are attached to clusters of cells in the population. These observations, combined with the established role of exoenzymes in the formation of MnO_X_ precipitates, suggests that these provide a community function to AzwK-3b and allows cultures grown to sufficient density in the presence of manganese to become resistant to the inhibitory effects of nitrite. Our attempts to elucidate the mechanistic basis of this functionality showed that biogenic MnO_X_ can oxidise nitrite to nitrate (under conditions that synthetic MnO_2_ cannot). Together with the known ability of MnO_X_to degrade hydrogen peroxide (40, 46–53), these findings show that biogenic MnO_X_ can inhibit the two key routes to the formation of reactive nitrite species.

These findings provide for the first-time a direct evidence for the impact of MnO_X_ on an organism’s growth, thus demonstrating a positive fitness effect and a possible evolutionary explanation to the costly process of MnO_X_ oxidation. Other suggested functional roles for this process to date were either hypothetical or were based on experiments with synthetic manganese oxide counterparts (2, 7, 8), and none of them were fully confirmed in a biological context. While mitigation of nitrite inhibition might not be the only evolutionary advantage of MnO_X_ oxidation in AzwK-3b or other manganese oxidizing species, it is definitely an ecologically relevant function. Nitrite is a known inhibitor in the environment (37, 38, 60), including in wastewater treatment applications (39). In the case of AzwK-3b, this ecological relevance is highly suggestive, as this species was isolated from an "agriculturally impacted, shallow salt marsh" (16) where nitrite (among other nitrogen species) can occur due to microbial conversion of nitrogen fertilizers (61–64). It is also interesting to note that oceanic manganese-rich modules are found to contain both manganese oxidizing and reducing bacteria (6), with current-day representatives of the latter group, such as *Shewenalla oneidensis* (9), also being nitrate-reducers (65–67).

Our study opens up additional investigations into the mechanism of nitrite toxicity and the role of MnO_X_ oxidation in it. Multiple mechanisms of nitrite-inhibition of bacteria have been reported (37, 38), and a key role for free nitrous acid (i.e. protonated nitrite) (39) and peroxynitrite, from nitrite and hydrogen peroxide (40–42), is proposed. Both molecules can prevent chemiosmotic coupling, and are primarily formed at low pH (nitrite is often found to inhibit bacteria growth at pH < 7 (41, 42)). The formation of these reactive nitrite species can be enhanced in the vicinity of the cells, where a locally lowered pH (from chemiosmotic coupling) and an increased hydrogen peroxide concentration (due to cellular metabolic activity (44, 45, 54, 58, 68–75)) can be formed. Interestingly, these very local conditions could be avoided through the presence of MnO_X_, which can degrade hydrogen peroxide and catalyse the oxidation of nitrite to nitrate, which is a proton consuming process with increased rate at low pH (43). The latter proposition is confirmed here for the first time, as we show that biogenic MnO_X_ can catalyze nitrite oxidation even under physiological conditions (i.e. pH 8).

The former hypothesis, i.e. that MnO_x_ can interfere with nitrite toxicity operating through peroxynitrite formation with hydrogen peroxide remains to be fully confirmed. Our experiments with spikes of hydrogen peroxide did not alter the gross dynamics of MnO_X_ mediated nitrite-inhibition relief, but this could be due to the design of these experiments with hydrogen peroxide delivered in single doses rather than being delivered in a controlled manner in the vicinity of the cells. A single dose could have been readily dealt with additional peroxidases, without altering MnO_X_ mediated effects. On the other hand, our observation that the nitrite-stress is fully mitigated in NADH-supplemented cultures (even in the absence of MnO_X_) lends support to the idea that nitrite stress is mediated primarily through formation of peroxynitrite. In that case, the reductive power of NADH could be employed by peroxidases, as well as MnO_X_, to reduce hydrogen peroxide (54–56) and thereby stopping the formation of peroxynitrite, explaining the observed mitigation effect of NADH.

These possible mechanistic scenarios of nitrite toxicity and roles of NADH, peroxidases, and MnO_X_ in mitigating it, raise the question about why cells that already have several peroxidases, such as AzwK-3b (17, 59) (see also Table S1), might invest additional energy in the formation of MnO_X_ precipitates. The answer might relate to the exact reaction mechanisms of ROS scavenging. It has been suggested, for example, that different ROS scavenging enzymes have different substrate affinities and efficiencies (58). In this context MnO_X_ – mediated scavenging could be preferred under certain ROS concentrations and modes of production. In addition, and unlike peroxidases that require stoichiometric equivalents of reductans as e.g. NADH/NADPH for hydrogen peroxide reduction (57, 58), MnO_x_ at its different oxidation states (II, III, IV) can directly catalyze degradation of hydrogen peroxide without NADH involvement (18, 19, 21, 40, 46–53). The fact that some peroxidases, as well as the AzwK-3b enzyme catalyzing MnO_X_ formation, are exoenzymes (17, 76) could be also highly relevant. The expression of such exoenzymes is a ‘social trait’, that can be exploited by cheating cells that do not invest the costs but reap the benefits (32–35). The presented finding that MnO_X_ forms dispersed granules in the culture shows that, in this case, the functional effects of the exoenzyme is localized. This kind of localization is a known strategy to stabilize a social trait in the face of evolution of cheating, as seen in exoenzymes with localized actions, involved in sugar degradation (77) and metal scavenging (78). Thus, the NADH investment into the formation of MnO_x_ mediated protection might be a metabolically less costly strategy that is also socially more stable.

Within a wider context, our findings could be highly relevant to understand the different forms of metal mineralization observed in different microorganisms and under different ecological contexts. Given the abundance of microorganisms being involved in reactions of the nitrogen cycle, there is indeed potential transient accumulation of nitrite in different environments. It is also possible that MnO_X_ (or other minerals) can provide more broad protection against ROS chemistry. For example, manganese oxidation is also observed in spore-forming bacteria (79, 80), fungi and other microorganisms (as reviewed and shown in (2, 36)), where a role for nitrite stress remains to be elucidated. Our findings will facilitate such further studies of bio-mineralizing organisms and their different functional motives and social strategies.

## Materials and Methods

### Bacterial Strain and Culture Conditions

*Roseobacter sp.* AzwK-3b was obtained from Colleen Hansel (Woods Hole Oceanographic Institution, Falmouth, MA/USA), who isolated the strain (16). Cultures were grown in a defined medium, which was established by modifying the pre-defined artificial seawater (ASW) medium (22). This media is referred to as ASW_m_ from now on, and its composition is shown in Table 1. ASW_m_ contained sodium acetate as the sole carbon source (at concentrations specified per experiment), 200mM sodium chloride (instead of 428 mM, as in ASW), ammonium as nitrogen source (instead of nitrate, as in ASW), and five vitamins that were added as supplement. In manganese-supplemented ASW_m_, manganese chloride (MnCl_2_) was added to 200 μM. Cultures were grown at 30 °C in appropriate (100 ml) Erlenmeyer flasks (shaking at 150 rpm) or 96 well polystyrene plates (Corning Inc.) closed with lid and parafilm (shaking at 300 rpm). Plates were incubated in a CLARIOstar plate reader (BMG labtech) and absorbance measurements were done at 600 nm (A_600_) and with path length-correction, so to present absorbance per 1 cm.

### Electron microscopy (EM) and Energy Dispersive X-ray spectroscopy (EDS) analysis of AzwK-3b cultures

A culture of AzwK-3b (40 ml in 100 ml Erlenmeyer flasks) was inoculated in ASW_m_ without manganese and nitrite, and containing 50 mM acetate. After 3 days at 150 rpm and 30 °C (by which time the culture reached the stationary phase), dilutions (25x – 200x) were made for a second passage of culture in the same medium, supplemented with 200 μM manganese. After further 2 days of culturing, samples for EM were prepared as follows: Cells from 2.5 ml culture were harvested by centrifugation (5 min at 5,000 g), and the supernatant was discarded. From here, several washing and dehydration steps were conducted by resuspending the pellet in different solutions and subsequently centrifuging for 5 min at 5,000 g (supernatant discarded): (1) first, pellets were twice re-suspended in ASW_m_ medium basis (no manganese, no acetate, no ammonium, no nitrite, no trace metals); (2) afterwards, samples were re-suspended in 200 μl 70 % ethanol, incubated for 1 min, and pelleted by centrifugation; (3) for a washing-dehydration step, pellets were twice re-suspended in 200 μl 100 % ethanol and harvested by centrifugation; (4) finally, samples were re-suspended in 100 μl of 100 % ethanol. This suspension was then applied to Transmission Electron Microscopy (TEM) grids (Lacey carbon film coated copper grids (Agar Scientific)) by pipetting, in 1 μl portions (allowed to dry in between), until a total of 2 or 5 μl was accumulated (on different grids).

EM analysis was done on a Gemini SEM 500 (Zeiss) equipped with EDS X-Max detector (Oxford Instruments). Data analysis was done on the associated AZtec software, which contained the spectral information to identify individual elements. Electron micrographs had the best quality in scanning transmission EM mode (STEM) with a high angle annular dark field detector (HAADF). For EDS, the sample needed to be moved, and the HAADF detector had to be withdrawn, so the location of analysis after changing the setup was confirmed by additional scanning EM (SEM) recording.

### Large inocula preparation for nitrite-assays

AzwK-3b was grown in Erlenmeyer flasks (usually 40 ml culture volume in 100 ml Erlenmeyer flasks) in ASW_m_ with 25 mM acetate. The culture absorbance A_600_ was recorded regularly on a Spectronic 200 spectrophotometer (Thermo Fisher) with 1 cm path length polystyrene cuvettes, and inocula were sampled at various stages of the growth curve (e.g. see Figures S4, S5, S8). This culture was used to inoculate into 96 well plates, which were supplemented by 1:1 dilution with fresh medium supplemented with manganese and/or nitrite and other additives, as described for the particular results shown (see legends of Figures 3, 7, S6, S8). Where noted (see respective figure captions), the fresh medium used for dilution was also supplemented with NADH or hydrogen peroxide at different concentrations. NADH or hydrogen peroxide were added as last additives (to prevent reaction e.g. between hydrogen peroxide and Mn^II^ before inoculation) and the completed fresh medium was used immediately.

### Growth curve fitting and analysis

Growth curves were analyzed using the R-package Grofit (23) applying the Gompertz growth model (23, 24). Plate reader data (measurements every 10 minutes) were de-noised by averaging over 6 measurements (i.e. hourly averages). The maximum A_600_ reached was read directly from the data. For curve fitting, all data later than the maximum A_600_, i.e. decaying growth phase, were removed. Then, the data was read backwards in time to find the first reading that was below 5 % of the maximum A_600_. This data-trimming was done to facilitate the fitting of the Gompertz growth model without bias from different lag-phases (which were ignored), or different lengths and scales of decaying phases recorded. From the resulting model, the maximum growth rate μ (in A_600 nm_(a.u.) per hour) was recorded.

### Preparation of cell-free bio-manganese oxide

The procedure was adapted from previous publications using the cell free supernatant of *Roseobacter sp.* AzwK-3b grown in complex medium (16–19). AzwK-3b was grown in ASW_m_ supplemented with 50 mM sodium acetate for nine days, using individual 50 or 100 ml cultures in 100 or 200 ml Erlenmeyer flasks, respectively, at 30 °C with shaking (150 rpm). In total, 2 liters of culture was prepared, cells were removed by centrifugation (5 minutes at 10,000 g) and the supernatants were combined. From this (cell-free) supernatant, individual samples of 100 or 200 ml were prepared and supplemented with 200 μM manganese chloride, MnCl_2_. Manganese oxidation was allowed to proceed for five days at 30 °C with shaking (150 rpm), after which the manganese oxide was harvested by centrifugation (5 minutes at 10,000 g) from each 50/100 ml sample. These were combined and washed by suspending in 25 ml acetate-free ASW_m_ medium and re-sedimented by centrifugation. The pellet was brown in appearance and had considerable volume, indicating co-precipitation of organic material (e.g. secreted proteins) from the cell-culture supernatant. To estimate the amount of manganese precipitated in the assay, the supernatants from centrifugation and the washing steps were combined, and the residual manganese determined by the 3,3’,5,5’-tetramethylbenzidine (TMB)-assay (25) for soluble manganese. Note that this was not a precise quantification, but was conclusive enough to allow conservative stoichiometric relations to be inferred. In particular, we inferred that ca. 75 % of the 200 μM manganese chloride had been removed from the solution and this value was used for downstream calculations. The MnO_X_ precipitate was suspended in an appropriate volume of the acetate-free medium to produce a “10 mM” suspension of manganese oxide, and this value is used in the manuscript as indicator for manganese oxide concentration. The pH was 8.2, which is well in line with the pH 8.0 of the ASW_m_ medium, showing that the suspended manganese oxide did not alter the pH.

### Quantification of nitrite, nitrate and acetate

Quantification was done by Ion Chromatography (IC) on a DIONEX ICS-5000+ (ThermoFisher, UK) equipped with conductivity detector and a DIONEX IonPac AS11-HC-4μm (2 × 250 mm ThermoFisher, UK) anion separation column with appropriate guard column. Separation was achieved with a potassium hydroxide (KOH) gradient, with the KOH added to the eluent by electrolytic eluent generation and, before conductivity detection, removed by electrochemical eluent suppression (both the generation and suppression units are part of the ICS-5000+ system). Culture samples were filtered (0.22 μm polyamide spin filter Costar Spin-X, Corning, NY/USA) and 10-fold diluted with MilliQ-water (checked for purity by measuring resistance (R); R > 18.2 MΩ), of which 2.5 μl were injected for IC separation. The IC was run at flow rate of 0.38 ml/min, column temperature 30 °C, and a conductivity detector cell temperature of 35 °C. The gradient condition, for the 37 minutes total run-time including 7 minutes pre-equilibration time, was: 7 minutes pre-run (equilibration) at 1.5 mM KOH before injection; remain 8 minutes at 1.5 mM KOH; increase to 15 mM KOH over 10 minutes; increase to 24 mM KOH over 5 minutes; increase to 60 mM KOH over 1 minutes; remain at 60 mM KOH over 6 minutes. Reference samples with known concentrations were run for calibration, from which the concentrations of nitrite, nitrate, and acetate in the samples was inferred. During the course of the experiments (see below) evaporation of the samples was noted (indicated by the increase in the peak area of chloride, which is expected to be unaltered by any biologic means and therefore should have displayed no concentration change). To correct for this evaporation effect, the concentrations of the analytes of interest were corrected by the same ratio as that obtained from the chloride peak area (from the beginning and end point samples of a particular time-course experiment).

## Author Contributions

CZ, JCO and OSS designed the study and the experiments. CZ performed the experiments and analyzed the data. All authors contributed to the writing of the manuscript and have given approval to the final version.

## Acknowledgments

This work is funded by The University of Warwick and by the Biotechnological and Biological, Natural Environment, and Engineering and Physical Sciences Research Councils (BB–, NE-, and EPSRC), with grant IDs: BB/K003240/2 (to OSS), NE/K009044/1 (to JCO) and BB/M017982/1 (to the Warwick Integrative Synthetic Biology Centre, WISB). We would like to thank Colleen Hansel (Woods Hole Oceanographic Institution) for providing *Roseobacter sp.* AzwK-3b, and Steve York from the Electron Microscopy research technology platform (EM RTP) at the Materials Science Department (Physics, University of Warwick) for EM/EDS measurements.

## References

1. Lowenstam H (1981) Minerals formed by organisms. Science 211(4487):1126–1131.

2. Hansel CM (2017) Manganese in marine microbiology. Advances in Microbial Physiology - Microbiology of Metal Ions, ed Poole RK (Academic Press, Oxford), pp 37–83.

3. Nealson KH (2006) The manganese-oxidizing bacteria. Prokaryotes 5:222–231.

4. Ghiorse WC (1984) Biology of iron-and manganese-depositing bacteria. Annu Rev Microbiol 38(1):515–550.

5. Spiro TG, Bargar JR, Sposito G, Tebo BM (2010) Bacteriogenic manganese oxides. Acc Chem Res 43(1):2–9.

6. Blöthe M, Wegorzewski A, Müller C, Simon F, Kuhn T, Schippers A (2015) Manganese-cycling microbial communities inside deep-sea manganese nodules. Environ Sci Technol 49(13):7692–7700.

7. Tebo BM, Johnson HA, McCarthy JK, Templeton AS (2005) Geomicrobiology of manganese(II) oxidation. Trends Microbiol 13(9):421–428.

8. Geszvain K, Butterfield C, Davis RE, Madison AS, Lee S-W, Parker DL, Soldatova A, Spiro TG, Luther GW, Tebo BM (2012) The molecular biogeochemistry of manganese(II) oxidation. Biochem Soc Trans 40(6):1244–1248.

9. Myers CR, Nealson KH (1988) Bacterial manganese reduction and growth with manganese oxide as the sole electron acceptor. Science 240(4857):1319–21.

10. Venkateswaran K, Moser DP, Dollhopf ME, Lies DP, Saffarini DA, MacGregor BJ, Ringelberg DB, White DC, Nishijima M, Sano H, Burghardt J, Stackebrandt E, Nealson KH (1999) Polyphasic taxonomy of the genus *Shewanella* and description of *Shewanella oneidensis sp. nov*. Int J Syst Bacteriol 49(2):705–24.

11. Lovley DR (1993) Dissimilatory metal reduction. Annu Rev Microbiol 47:263–90.

12. Emerson D, Fleming EJ, McBeth JM (2010) Iron-oxidizing bacteria: an environmental and genomic perspective. Annu Rev Microbiol 64:561–83.

13. Remucal CK, Ginder-Vogel M (2014) A critical review of the reactivity of manganese oxides with organic contaminants. Environ Sci Process Impacts 16(6):1247–66.

14. Sunda WG, Kieber DJ (1994) Oxidation of humic substances by manganese oxides yields low-molecular-weight organic substrates. Nature 367(6458):62–64.

15. Keiluweit M, Nico P, Harmon ME, Mao J, Pett-Ridge J, Kleber M (2015) Long-term litter decomposition controlled by manganese redox cycling. Proc Natl Acad Sci 112(38):E5253–E5260.

16. Hansel CM, Francis CA (2006) Coupled photochemical and enzymatic Mn(II) oxidation pathways of a planktonic *Roseobacter*-Like bacterium. Appl Environ Microbiol 72(5):3543–9.

17. Andeer PF, Learman DR, McIlvin M, Dunn JA, Hansel CM (2015) Extracellular haem peroxidases mediate Mn(II) oxidation in a marine *Roseobacter* bacterium via superoxide production. Environ Microbiol 17(10):3925–3936.

18. Learman DR, Voelker BM, Vazquez-Rodriguez AI, Hansel CM (2011) Formation of manganese oxides by bacterially generated superoxide. Nat Geosci 4(2):95–98.

19. Learman DR, Voelker BM, Madden AS, Hansel CM (2013) Constraints on superoxide mediated formation of manganese oxides. Front Microbiol 4:262.

20. Learman DR, Wankel SD, Webb SM, Martinez N, Madden AS, Hansel CM (2011) Coupled biotic–abiotic Mn(II) oxidation pathway mediates the formation and structural evolution of biogenic Mn oxides. Geochim Cosmochim Acta 75(20):6048–6063.

21. Luther GW (2010) The role of one- and two-electron transfer reactions in forming thermodynamically unstable intermediates as barriers in multi-electron redox reactions. Aquat Geochemistry 16(3):395–420.

22. Wilson WH, Carr NG, Mann NH (1996) The effect of phosphate status on the kinetics of cyanophage infection in the oceanic cyanobacterium *Synechococcus sp.* WH78031. J Phycol 32(4):506–516.

23. Kahm M, Hasenbrink G, Lichtenberg-Fraté H, Ludwig J, Kschischo M (2010) Grofit: fitting biological growth curves with R. J Stat Softw 33(7):1–21.

24. Zwietering MH, Jongenburger I, Rombouts FM, van ’t Riet K (1990) Modeling of the bacterial growth curve. Appl Environ Microbiol 56(6):1875–81.

25. Bosch Serrat F (1998) 3,3’,5,5’-Tetramethylbenzidine for the colorimetric determination of manganese in water. Mikrochim Acta 129(1–2):77–80.

26. Tebo BM, Bargar JR, Clement BG, Dick GJ, Murray KJ, Parker D, Verity R, Webb SM (2004) Biogenic manganese oxides: properties and mechanisms of formation. Annu Rev Earth Planet Sci 32(1):287–328.

27. Templeton AS, Staudigel H, Tebo BM (2005) Diverse Mn(II)-oxidizing bacteria isolated from submarine basalts at Loihi seamount. Geomicrobiol J 22(3–4):127–139.

28. Francis CA, Co E-M, Tebo BM (2001) Enzymatic manganese(II) oxidation by a marine α-proteobacterium. Appl Environ Microbiol 67(9):4024–4029.

29. Johnson HA, Tebo BM (2008) In vitro studies indicate a quinone is involved in bacterial Mn(II) oxidation. Arch Microbiol 189(1):59–69.

30. Wang X, Wiens M, Divekar M, Grebenjuk VA, Schröder HC, Batel R, Müller WEG (2010) Isolation and characterization of a Mn(II)-oxidizing *Bacillus* strain from the demosponge *Suberites domuncula*. Mar Drugs 9(1):1–28.

31. Kraft B, Strous M, Tegetmeyer HE (2011) Microbial nitrate respiration – Genes, enzymes and environmental distribution. J Biotechnol 155(1):104–117.

32. Cavaliere M, Feng S, Soyer OS, Jiménez JI (2017) Cooperation in microbial communities and their biotechnological applications. Environ Microbiol 19(8):2949–2963.

33. Allen B, Gore J, Nowak MA (2013) Spatial dilemmas of diffusible public goods. Elife 2:e01169.

34. West SA, Diggle SP, Buckling A, Gardner A, Griffin AS (2007) The social lives of microbes. Annu Rev Ecol Evol Syst 38(1):53–77.

35. Allison SD (2005) Cheaters, diffusion and nutrients constrain decomposition by microbial enzymes in spatially structured environments. Ecol Lett 8(6):626–635.

36. Keim CN, Nalini HA, de Lena JC (2015) Manganese oxide biominerals from freshwater environments in Quadrilatero Ferrifero, Minas Gerais, Brazil. Geomicrobiol J 32(6):549–559.

37. Cammack R, Joannou C., Cui X-Y, Torres Martinez C, Maraj SR, Hughes MN (1999) Nitrite and nitrosyl compounds in food preservation. Biochim Biophys Acta - Bioenerg 1411(2–3):475–488.

38. Müller-Herbst S, Mühlig A, Kabisch J, Rohtraud Pichner, Scherer S (2015) The food additives nitrite and nitrate and microbiological safety of food products. Am J Microbiol 6(1):1–3.

39. Zhou Y, Oehmen A, Lim M, Vadivelu V, Ng WJ (2011) The role of nitrite and free nitrous acid (FNA) in wastewater treatment plants. Water Res 45(15):4672–82.

40. Robinson KM, Beckman JS (2005) Synthesis of peroxynitrite from nitrite and hydrogen peroxide. Methods Enzymol:207–214.

41. Heaselgrave W, Andrew PW, Kilvington S (2010) Acidified nitrite enhances hydrogen peroxide disinfection of acanthamoeba, bacteria and fungi. J Antimicrob Chemother 65(6):1207–14.

42. Kono Y, Shibata H, Adachi K, Tanaka K (1994) Lactate-dependent killing of *Escherichia coli* by nitrite plus hydrogen peroxide: a possible role of nitrogen dioxide. Arch Biochem Biophys 311(1):153–159.

43. Luther, III GW, Popp JI (2002) Kinetics of the abiotic reduction of polymeric manganese dioxide by nitrite: an anaerobic nitrification reaction. Aquat Geochemistry 8(1):15–36.

44. Martínez MC, Andriantsitohaina R (2009) Reactive nitrogen species: molecular mechanisms and potential significance in health and disease. Antioxid Redox Signal 11(3):669–702.

45. Tharmalingam S, Alhasawi A, Appanna VP, Lemire J, Appanna VD (2017) Reactive nitrogen species (RNS)-resistant microbes: adaptation and medical implications. Biol Chem 398(11):1193–1208.

46. Watts RJ, Sarasa J, Loge FJ, Teel AL (2005) Oxidative and reductive pathways in manganese-catalyzed Fenton’s reactions. J Environ Eng 131(1):158–164.

47. Jiang S, Ashton WR, Tseung ACC (1991) An observation of homogeneous and heterogeneous catalysis processes in the decomposition of H_2_O_2_ over MnO_2_ and Mn(OH)_2_. J Catal 131(1):88–93.

48. Kanungo SB, Parida KM, Sant BR (1981) Studies on MnO_2_—III. The kinetics and the mechanism for the catalytic decomposition of H_2_O_2_ over different crystalline modifications of MnO_2_. Electrochim Acta 26(8):1157–1167.

49. Rophael MW, Petro NS, Khalil LB (1988) II — kinetics of the catalytic decomposition of hydrogen peroxide solution by manganese dioxide samples. J Power Sources 22(2):149–161.

50. Do S-H, Batchelor B, Lee H-K, Kong S-H (2009) Hydrogen peroxide decomposition on manganese oxide (pyrolusite): Kinetics, intermediates, and mechanism. Chemosphere 75(1):8–12.

51. Li W, Liu Z, Liu C, Guan Y, Ren J, Qu X (2017) Manganese dioxide nanozymes as responsive cytoprotective shells for individual living cell encapsulation. Angew Chem Int Ed Engl 56(44):13661–13665.

52. Broughton DB, Wentworth RL (1947) Mechanism of decomposition of hydrogen peroxide solutions with manganese dioxide. I. J Am Chem Soc 69(4):741–744.

53. Broughton DB, Wentworth RL, Laing ME (1947) Mechanism of decomposition of hydrogen peroxide solutions with manganese dioxide. II. J Am Chem Soc 69(4):744–747.

54. Seaver LC, Imlay JA (2001) Alkyl hydroperoxide reductase is the primary scavenger of endogenous hydrogen peroxide in *Escherichia coli*. J Bacteriol 183(24):7173–7181.

55. Farr SB, Kogoma T (1991) Oxidative stress responses in *Escherichia coli* and *Salmonella typhimurium*. Microbiol Rev 55(4):561–85.

56. Nathan C, Bryk R, Griffin P (2000) Peroxynitrite reductase activity of bacterial peroxiredoxins. Nature 407(6801):211–215.

57. Zamocky M, Furtmüller PG, Obinger C (2008) Evolution of catalases from bacteria to humans. Antioxid Redox Signal 10(9):1527–1548.

58. Mishra S, Imlay J (2012) Why do bacteria use so many enzymes to scavenge hydrogen peroxide? Arch Biochem Biophys 525(2):145–60.

59. Diaz JM, Hansel CM, Voelker BM, Mendes CM, Andeer PF, Zhang T (2013) Widespread production of extracellular superoxide by heterotrophic bacteria. Science 340(6137):1223–6.

60. Camargo JA, Alonso A (2006) Ecological and toxicological effects of inorganic nitrogen pollution in aquatic ecosystems: A global assessment. Environ Int 32(6):831–49.

61. Cleemput O, Samater AH (1995) Nitrite in soils: accumulation and role in the formation of gaseous N compounds. Fertil Res 45(1):81–89.

62. Riley WJ, Ortiz-Monasterio I, Matson PA (2001) Nitrogen leaching and soil nitrate, nitrite, and ammonium levels under irrigated wheat in Northern Mexico. Nutr Cycl Agroecosystems 61(3):223–236.

63. Lawniczak AE, Zbierska J, Nowak B, Achtenberg K, Grześkowiak A, Kanas K (2016) Impact of agriculture and land use on nitrate contamination in groundwater and running waters in central-west Poland. Environ Monit Assess 188(3):172.

64. Beeckman F, Motte H, Beeckman T (2018) Nitrification in agricultural soils: impact, actors and mitigation. Curr Opin Biotechnol 50:166–173.

65. Cruz-García C, Murray AE, Klappenbach JA, Stewart V, Tiedje JM (2007) Respiratory nitrate ammonification by *Shewanella oneidensis* MR-1. J Bacteriol 189(2):656–662.

66. Chen Y, Wang F (2015) Insights on nitrate respiration by *Shewanella*. Front Mar Sci 1:80.

67. Zhang H, Fu H, Wang J, Sun L, Jiang Y, Zhang L, Gao H (2013) Impacts of Nitrate and Nitrite on Physiology of *Shewanella oneidensis*. PLoS One 8(4):e62629.

68. Quijano C, Trujillo M, Castro L, Trostchansky A (2016) Interplay between oxidant species and energy metabolism. Redox Biol 8:28–42.

69. Davies KJ (1995) Oxidative stress: the paradox of aerobic life. Biochem Soc Symp 61:1–31.

70. Gutteridge JM (1994) Biological origin of free radicals, and mechanisms of antioxidant protection. Chem Biol Interact 91(2–3):133–40.

71. Korshunov S, Imlay JA (2010) Two sources of endogenous hydrogen peroxide in *Escherichia coli*. Mol Microbiol 75(6):1389–1401.

72. van der Heijden J, Vogt SL, Reynolds LA, Peña-Díaz J, Tupin A, Aussel L, Finlay BB (2016) Analysis of bacterial survival after exposure to reactive oxygen species or antibiotics. Data Br 7:894–899.

73. van der Heijden J, Vogt SL, Reynolds LA, Peña-Díaz J, Tupin A, Aussel L, Finlay BB (2016) Exploring the redox balance inside gram-negative bacteria with redox-sensitive GFP. Free Radic Biol Med 91:34–44.

74. Seaver LC, Imlay JA (2001) Hydrogen peroxide fluxes and compartmentalization inside growing *Escherichia coli*. J Bacteriol 183(24):7182–7189.

75. González-Flecha B, Demple B (1995) Metabolic sources of hydrogen peroxide in aerobically growing *Escherichia coli*. J Biol Chem 270(23):13681–13687.

76. Christie-Oleza JA, Scanlan DJ, Armengaud J (2015) “You produce while I clean up”, a strategy revealed by exoproteomics during *Synechococcus* - *Roseobacter* interactions. Proteomics 15(20):3454–3462.

77. Gore J, Youk H, van Oudenaarden A (2009) Snowdrift game dynamics and facultative cheating in yeast. Nature 459(7244):253–6.

78. Kümmerli R, Jiricny N, Clarke LS, West SA, Griffin AS (2009) Phenotypic plasticity of a cooperative behaviour in bacteria. J Evol Biol 22(3):589–98.

79. Francis CA, Tebo BM (2002) Enzymatic manganese(II) oxidation by metabolically dormant spores of diverse *Bacillus* species. Appl Environ Microbiol 68(2):874–880.

80. Bargar J., Tebo B., Villinski J. (2000) In situ characterization of Mn(II) oxidation by spores of the marine *Bacillus sp.* strain SG-1. Geochim Cosmochim Acta 64(16):2775–2778.

81. Tang YJ, Laidlaw D, Gani K, Keasling JD (2006) Evaluation of the effects of various culture conditions on Cr(VI) reduction by *Shewanella oneidensis* MR-1 in a novel high-throughput mini-bioreactor. Biotechnol Bioeng 95(1):176–184.

82. Balch WE, Fox GE, Magrum LJ, Woese CR, Wolfe RS (1979) Methanogens: reevaluation of a unique biological group. Microbiol Rev 43(2):260–96.

